# The deubiquitinating enzyme CYLD impairs NF-κB- and STAT1-dependent macrophage intrinsic immunity to *Staphylococcus aureus*

**DOI:** 10.1101/2024.07.25.605106

**Authors:** Christina Schmidt, Kunjan Harit, Stephan Traidl, Thomas Werfel, Lennart Rösner, Gopala Nishanth, Dirk Schlüter

**Author notes:** These authors share senior authorship (Lennart Rösner, Gopala Nishanth, Dirk Schlüter).

## Abstract

In atopic dermatitis (AD), lesional skin is frequently colonized by *Staphylococcus* (*S*.) *aureus*, contributing to the severity and clinical symptoms of the disease. The inflammatory milieu in the skin is characterized by a type 2 inflammatory signature, including M2 macrophages, which cannot eradicate S. aureus. Since S. aureus is effectively controlled in macrophages activated by pattern recognition receptor (PRR)-induced NF-κB and interferon (IFN)-γ-induced STAT1 stimulation, we hypothesized that pre-treatment with LPS as a PRR/TLR4-activating agent and IFN-γ would induce effective control of S. aureus in monocyte-derived macrophages (MDMs) of AD patients. Our data show that the deubiquitinating enzyme CYLD is strongly expressed in skin macrophages and MDMs from AD patients compared to healthy controls and impairs the anti-staphylococcal activity of PRR-activated and IFN-treated MDMs. Functionally, CYLD impaired M1 macrophage polarization, as evidenced by reduced expression of CD80, TNF, and IL-6 upon LPS- and IFN-γ treatment in CYLD-deficient as compared to wild-type (WT) MDM/THP1 macrophages. Mechanistically, CYLD inhibited IFN-γ-induced STAT1 phosphorylation and activation by binding to STAT1 and inducing its K63 deubiquitination. CYLD also interacted with TRAF6 and NEMO/IKKγ in the MYD88 signaling pathway and with RIPK2 in the NOD2 pathway, leading to impaired activation of NF-κB. Inhibition of both STAT1 by siRNA and NF-κB by IKK inhibitor treatment, respectively, independently abolished the control of *S. aureus* in both CYLD-deficient and WT THP1 macrophages, which harbored identical high numbers of the pathogen. The in vivo inhibitory function of CYLD on the control of *S. aureus* was also observed upon infection of *Cyld*-deficient and WT mice. Collectively, these data illustrate that the increased expression of CYLD in macrophages of AD patients is a factor contributing to the ineffective control of *S. aureus*, diminishes M1 macrophage polarization, and that CYLD deletion unleashes the break on effective STAT1 and NF-κB-dependent control of *S. aureus*.

## 1 Introduction

Eczematous skin lesions of patients with atopic dermatitis (AD) are frequently colonized with *Staphylococcus* (*S*.) *aureus* and the extent of colonization increases in more severe lesions with hyperinflammation (Edslev et al., 2021; Guzik et al., 2006; Kong et al., 2012; Saheb Kashaf et al., 2023; Wichmann et al., 2009). These staphylococci contribute to disease symptoms, in particular itching and pain, by the secretion of toxins and proteases which directly stimulate sensory nerves in the skin (Deng et al., 2023; Gheoghegan et al., 2018; Langer et al., 2007). In addition, these staphylococci may cause local and, in some cases, systemic infections in AD patients. The importance of the role of *S. aureus* in AD is supported by the fact that treatment of the skin infection with antiseptics or antibiotics can improve the symptoms of AD (Breuer et al., 2002).

Upon infection, control of *S. aureus* critically depends on protective innate immune responses. Neutrophils can phagocytose and kill the bacteria and also immobilize *S. aureus* through NETosis, i.e. capturing the bacteria in a dense network of DNA released by the neutrophils (Spaan et al., 2013; von Köckritz-Blickwede and Winstel; 2022)). In addition, both tissue-resident and monocyte-derived macrophages contribute to the control and elimination of *S. aureus* (Pidwill et al., 2021). Macrophages can rapidly phagocytose *S. aureus* and activate the NF-κB pathway which contributes to the production of anti-bacterial reactive oxygen-species (ROS) and nitric oxide (Surewaard et al., 2016). Activation of macrophages by interferon (IFN)-γ synergizes with TLR2-mediated NF-κB activation in NO production and further enhances control of *S. aureus* in macrophages (citation). These “M1-like” macrophages also contribute to the control of local *S. aureus* skin infection and contribute to the prevention of systemic bacterial dissemination (Chan et al., 2018). In contrast to M1 macrophages, M2 polarized macrophages have an impaired anti-staphylococcal activity and capacity to control *S. aureus* infections. (Pidwill et al., 2021). *S. aureus* can subvert these protective intracellular mechanisms to replicate and persist intracellularly in macrophages and other cell types including keratinocytes (Gresham et al., 2000; Soong et al., 2015; Surewaard et al., 2016; Thammavongsa et al., 2015; Tuchscherr et al., 2011).

The lesional skin of AD patients is characterized by a type-2 inflammatory milieu composed of by macrophages, monocytes, B cells, CD4^+^ and CD8^+^ T cells (Beck et al., 2022; Zhang et al., 2023). This type-2 inflammatory environment may facilitate persistence of extracellular *S. aureus* but also intracellularly in keratinocytes (Soong et. al., 2015) and in macrophages (Thammavongsa et al., 2015; Tuchscherr et al., 2011). In AD patients, the dysregulation of the immune system also extends beyond the skin and includes altered systemic innate and adaptive immune profile leading to comorbidities including infections (Brunner et al., 2017; Droitcourt et al., 2021; Jin et al., 2024).

Immune responses are critically regulated by post-translational modifications including ubiquitination. Ubiquitination is a highly dynamic process exerted by a cascade of ubiquitin-activating E1, ubiquitin-conjugating E2 and ubiquitin E3 ligases which can attach the ubiquitin protein to a lysine residue of a substrate protein and by deubiquitinating enzymes (DUBs) which can cleave ubiquitin from the substrates. Ubiquitin can be attached as monomers or as ubiquitin chains to the substrates. In polyubiquitination, ubiquitin molecules are linked through any one of the internal seven lysine (K6, K11, K27, K33, K48, K63) or the N-terminal methionine residue (M1). The type of ubiquitin linkage decides on the function and fate of the substrate and can lead to proteasomal degradation by K48- and branched K11/K48 linked ubiquitin chains (Chau et al., 1989; Meyer and Rape, 2014) or to K63-linked ubiquitin modification of protein function including regulation of signal transduction (Raman and Wolberger, 2024).

Ubiquitination is reversible and counteracted by DUBs. The DUB CYLD can cleave K63- and M1-linked polyubiquitin chains from substrates. and regulates a broad range of key cellular processes including inflammatory responses, cell death pathways, autophagy, DNA damage and cell proliferation (Komander et al., 2009; Marin-Rubio et al., 2023). In immune signaling, CYLD negatively regulates NF-κB by deubiquitinating TRAF2, TRAF6, TAK1, and NEMO (Brummelkamp et al., 2003; Kovalenko et al., 2003; Reiley et al., 2007; Trompouki et al., 2003). With respect to bacterial infections, this suppressive effect of CYLD on NF-κB prevented immunopathology in *Haemophilus influenzae* and *Streptococcus pneumoniae* infections (Lim et al., 2007a; Lim et al., 2007b) but protective immune responses in *Escherichia coli* pneumonia and in systemic listeriosis (Lim et al., 2008; Wurm et al., 2015). In addition, CYLD deubiquitinates STAT3 and RIPK2, thereby, impairing protective IL-6 and NOD2-dependent protective immunity in listeriosis (Nishanth et al., 2013; Wex et al., 2016).

Inactivating mutations of *CYLD* underlie the CYLD cutaneous syndrome, a disease characterized by the development of benign skin tumors of the hair follicles (Nagy et al., 2021) including multiple familial trichoepithelioma, the Brooke–Spiegler syndrome, and familial cylindromatosis. In addition, somatic *CYLD* mutations have been linked to suppression of sporadic cancers, non-alcoholic steatohepatitis (Ji et al., 2018)), arterial hypertension (Zhou et al., 2021) and by gain of function mutations to neurodegenerative disorders (Dobson-Stone et al., 2020) (reviewed in Marin-Rubio et al., 2023).

Although CYLD is strongly expressed in the healthy skin and in macrophages (Marin-Rubio et al., 2023; Uhlen et al., 2010; https://www.proteinatlas.org/ENSG00000083799-CYLD/single+cell+type), its expression and function in AD and *S. aureus* infections are unknown. To address these open questions, we analyzed the function of CYLD in AD patients. Our data show that CYLD is strongly expressed in the skin of AD patients and that CYLD impairs STAT1 and NF-κB-dependent control of *S. aureus* in macrophages.

## 2 Material and Methods

### Ethics statement

All animal experiments were in compliance with the German animal protection law in a protocol approved by the Landesverwaltungsamt Sachsen-Anhalt (file number: 203.h-42502-2-901, University of Magdeburg). The ethics committee of Hannover Medical School (MHH) approved the parts of the study involving patients (No. 10499-BO-K-2022).

### Animals

Age and sex matched animals were used for the experiments. C57BL/6 WT were obtained from Janvier (Le Genest Saint Isle, France) and C57BL/6 *Cyld*^-/-^ mice were kindly provided by Dr. Ramin Massoumi (Department of Laboratory Medicine, Malmö, Sweden) (Massoumi et al., 2006). All animals were kept under SPF conditions in an isolation facility of the Otto-von-Guericke University Magdeburg (Magdeburg, Germany).

### Patients

Patients with AD were recruited at the Department of Dermatology and Allergy of Hannover Medical School (MHH). The work described has been carried out in accordance with the Code of Ethics of the World Medical Association (Declaration of Helsinki), and patients gave their written informed consent prior to the study.

#### Staphylococcus aureus

Wild type (strain SH1000) and methicillin-resistant (strain MW2) *S. aureus* were grown in Luria broth (LB, Oxoid, Germany) and aliquots of log-phase cultures were stored at - 80°C. For infection of cells, fresh log phase cultures were prepared from frozen stock.

### THP-1 cells

The THP-1 cells (clone TIB-202) were obtained from the American Type Culture Collection (ATCC, Manassas, VA, USA). The cells were cultured in in RPMI cell culture medium supplemented with 10% fetal calf serum (FCS) and 1% penicillin/streptomycin.

### Generation of *CYLD*-deficient THP-1 cells

*CYLD* was knocked down by the CRISPR/Cas9 method using SG cell line 4D-nucleofector X kit (#V4XP-4024, Lonza, Basel, Switzerland). For 1x10^6^ cells, 210pmol of duplexed gRNA was mixed with 70pmol of Cas9 Nuclease V3 (#1081059, IDT, Iowa, USA) for RNP complex synthesis and the complex was mixed with 70pmol of EE buffer (#1075916, IDT) to form electroporation mix. Electroporation was performed using preset program DZ100 in Lonza nucleofection system.

### Generation of macrophages from THP-1 cells

THP-1 cells were differentiated into macrophages through M-CSF (50 ng/ml) treatment for 48 hours. The macrophages were either maintained in an unpolarized state (M0) or polarized into M1 phenotype by stimulation with IFN-γ (20 ng/ml) and LPS (10 pg/ml).

### Generation of monocyte-derived macrophages (MDM)

PBMCs were isolated using Ficoll density gradient centrifugation followed by magnetic cell separation of CD14^+^ monocytes (#480094, MojoSort, Biolegend, CA, USA). Stable knockout of *CYLD* was generated using CRISPR/Cas9 system and SG cell line 4D-nucleofector X kit (#V4XP-3024, Lonza) according to the manufactureŕs protocol. CD14^+^ monocytes were then cultured for 5 days in DMEM medium supplemented with 50ng M-CSF. Cells were harvested and stimulated as per experimental requirement.

### *In vitro* infection of cells with *S. aureus*

1x10^6^ THP-1 macrophages and MDM, respectively, were stimulated in 6-well plates (Greiner bio-one, Frickenhausen, Germany) with *S. aureus* at a multiplicity of infection (MOI) of 1:1 in RPMI supplemented with 10 % FCS. For infection, *S. aureus* was thawed from frozen stocks (-80 °C) and added to LB broth. The optical density of–log-phase cultures was determined using a photometer (Eppendorf, Hamburg, Germany). Cultures with an optical density of 0.1, which corresponds to a dose of 1x10^8^ *S. aureus*/ml according to a previously established standard growth curve, were pelleted by centrifugation (870 g, 4°C, 10 min) and the MOI was adjusted to 1:1 in LB supplemented with 10 % FCS. In each experiment, the bacterial dose used for infection was controlled by plating an inoculum on LB agar and counting colonies after incubation at 37 °C for 24h. After 1h of infection (37°C, 5% CO_2_), 30 µg/ml gentamicin (Sigma-Aldrich) was added for additional 30 min to kill extracellular bacteria. Thereafter, infected BMDM were washed twice with PBS and further cultivated in DMEM supplemented with gentamicin for the indicated time points. For inhibition of NF-κB, BMDM were treated with IKK inhibitor VII (1µM; Calbiochem, Darmstadt, Germany) starting 24 h before infection (concentration and incubation time were pretested before usage).

### CFUs

At the indicated time points p.i., *S aureus*-infected cells were centrifuged (5 min, 490 g) and the medium was discarded. Thereafter, the cells were washed with PBS to remove traces of remaining antibiotics. After centrifugation (5 min, 490 g) and removal of PBS, the cell pellets were lysed in 0.1% TritonX-100 and serial dilutions were plated on LB agar in petri dishes (ø 85 mm; Merck, Darmstadt, Germany). Bacterial colonies were counted microscopically using a grid plate after incubation at 37°C for 24 h and 48 h. The counting was conducted blindly and 16 fields were counted.

### Protein isolation and Western Blot

*S. aureus* infected THP-1 macrophages were washed in PBS and resuspended in 4-8 °C cold lysis buffer containing 50 mM Tris-HCl (pH 7.4), 5 mM EDTA, 100 mM NaCl, 1 % Triton-X-100, 10 % glycerol, 10 mM KH_2_PO_4_, 0.5% Na-deoxycholate, 1 mM phenylmethanesulfonyl fluoride 1 mM NaF, 1 mM Na_4_O_7_P_2_, 1 mM Na_3_VO_4_ and aprotinin, leupeptin, pepstatin (1µg/ml each) (all reagents from Sigma, Taufkirchen, Germany) for 30 min and centrifuged (19.000g, 4 °C, 10 min). The supernatant was collected and the protein concentration was determined by a commercial protein assay kit (Bio-Rad, Munich, Germany). A 1x lane marker reducing sample buffer (Thermo Scientific, Dreiech, Germany) was added to the samples and proteins were denatured at 99 °C for 5 min. Equal amounts of proteins were separated on SDS-polyacrylamide gels (6-12 %) and transferred semidry (220 mA, 60 min) on polyvinylidene fluoride membranes, preactivated in methanol. Unspecific binding of antibodies was blocked by incubating the membranes with Blotto B (1 % milkpowder plus 1 % bovine serum albumin (BSA), 5 % milkpowder and 5 % BSA, respectively, for 1h. The proteins were stained for GAPDH, phospho-STAT1 Y701, phospho-STAT1 S727, STAT-1, MyD88, IRAK-4, phospho-IRAK-4, TRAF6, NOD2, phospho-RIPK2, RIPK2, p65phospho-p65, p65, phospho-p38, p38, phospho-ERK1/2, ERK1/2, phospho-JNK, JNK, IKKγ/ NEMO, CYLD and K63-linkage-specific polyubiquitin (all antibodies from Cell Signaling Technology, Frankfurt, Germany), Primary antibodies were used at a dilution of 1:1000 in specific blocking medium as recommended by the supplier. Following overnight incubation, membranes were washed 3 times using Tris-Buffered Saline with 0.1%Tween 20 (TBST) for 10 minutes each and further incubated with 1:1000 diluted anti-mouse or anti-rabbit secondary antibodies (Dako, Hamburg, Germany) for 1 h. The blots were washed for 3 times in TBST for 10 minutes each and developed using an ECL Plus kit (GE Healthcare, Freiburg, Germany). For quantification of protein intensities by densitometry, WB images were captured using the Intas Chemo Cam Luminescent Image Analysis system^®^ (INTAS Science Imaging Instruments, Göttingen, Germany) and analyzed with the LabImage 1D software^®^ (Kapelan Bio-Imaging Solutions, Leipzig, Germany).

### Immunoprecipitation

Uninfected and *S aureus*-infected THP-1 macrophages as well as *CYLD*^-/-^ THP-1 macrophages were lysed on ice using the same protein lysis buffer as described above in “Protein isolation and Western Blot”. In a pre-clearing phase, Gamma Bind^TM^ G Sepharose^TM^ beads (GE Healthcare Europe GmbH, Freiburg Germany) were incubated with cell lysates under continuous shaking at 4°C for 30 min. The beads were removed by centrifugation (10 min, 10.000 g, 4°C) and equal amounts of lysates were incubated with anti-STAT1 (1:100) and anti-CYLD (1:100) antibodies, respectively, at 4°C overnight. The immune complexes were captured by incubating with fresh Gamma Bind^TM^ G Sepharose^TM^ beads at 4 °C overnight. The beads were then washed 3 times with PBS by pulse centrifugation (1000 g, 30 sec). The pellet containing the Gamma Bind^TM^ G Sepharose^TM^ immune complexes was resuspended in 1x lane marker reducing sample buffer and boiled at 99 °C for 5 min. Thereafter samples were centrifuged (1000 g, 30 sec), the supernatant was collected and used to detect STAT1, CYLD, K63, TRAF6, RIPK2, IKK/NEMO and K63-linked ubiquitin, by WB. GAPDH was used as the input control.

### Measurement of NO

The concentration of NO in the cell culture medium was measured using a Griess Assay Kit (Promega, Mannheim, Germany) according to the manufacturer’s instructions. In brief, the cells were centrifuged (400 g, for 5 min), to harvest the supernatant. Triplicates of diluted standard and supernatant (50 µl each) were added to the wells of a 96-well flat-bottom plate (Greiner bio-one, Frickenhausen, Germany). Subsequently, 50 µl of sulfanilamide solution was dispensed to the standard and the experimental samples and incubated in the dark for 10 min. Thereafter, 50 µl of *N*-(1-naphthyl) ethylenediamine solution was added to all samples, which were further incubated in the dark for 10 min. The NO concentration was determined by measuring samples at 540 nm using a Synergy^®^ microplate reader (Biotek, Berlin, Germany) within 30 min.

### ROS/RNI detection

The intracellular production of reactive oxygen species (ROS) and nitrogen species (RNI) was determined with a total ROS detection kit (Enzo Life Science, Lörrach, Germany). In brief, 1x10^6^ freshly prepared THP-1 macrophages were infected with *S. aureus* at an MOI of 1:1. After infection, the cells were centrifuged (400 g, for 5 min), the supernatant was discarded and the cell pellet was washed twice with 2 ml of 1x washing buffer. Thereafter, the cell pellet was resuspended in 500 µl detection solution (500µl 1x washing buffer + 0,1 µl detection reagent) and incubated in the dark at 37°C for 30 min. The samples were analyzed by Cytek Aurora flow cytometry (Fremont, CA).

### *In vitro* siRNA treatment

For siRNA-mediated knockdown of STAT1, WT and *CYLD*^-/-^ THP-1 macrophages were transfected with 20nm of STAT1 specific siRNA (Dharmacon, CO, USA) according to manufacturer’s instruction. 20µM of scrambled siRNA was used as control. A total of 50 µl of the siRNA mixture was added to each well in a 24-well plate containing 4x10^5^ macrophages and incubated at 37°C for 48 h. The efficiency of siRNA-mediated STAT1 silencing was controlled by WB.

### Immunostaining

OCT-embedded tissues were cut into 6-mm cryosections, air-dryed and fixed at 4°C in acetone at 4°C for 10min. After washing, HRP and AP blocking solution was applied for 20min (Invitrogen, Waltham, MA, USA), followed by Superblock (Thermo Scientific,Waltham, MA, USA). Rabbit polyclonal anti-CYLD (ab33929, Abcam, Cambridge, UK) or rabbit Ig (Dako X 0936, Agilent, Santa Clara, CA, USA) were applied in equal concentrations. Mouse monoclonal anti-CD3 (Agilent) was used to stain the cellular skin infiltrate. After applying the Envision+ HRP conjugate (Agilent) with anti-rabbit K4009 and anti-mouse K005, respectively, the AEC substrate kit was applied (AEC, 3-Amino-9-Ethylcarbazole; Zytomed Systems, Bargteheide, Germany). Images were taken on a Pannoramic MIDI II Sola (Sysmex, Norderstedt, Germany).

### Single cell RNA sequencing

Data from a previous study that included single-cell RNA sequencing were used for the investigation of CYLD expression on single cell level (Zhang et al., 2023). For a comprehensive explanation, refer to the corresponding methods section of the study. Briefly, skin punch biopsies were processed using the Miltenyi Biotech skin dissociation kit for subsequent cell isolation and CD2 enrichment, followed by pooling and loading onto the chromium chip (10x Genomics, Pleasanton, CA, USA). The CellRanger pipeline version 3.1.0 was employed to align reads to the human reference genome GRCH38. The expression matrix generated was then analyzed using the Seurat package version 4.2.3. UMAP and feature plots were generated using their respective functions.

### Statistics

The statistical significance was determined using software Prism 9 using respective tests as mentioned in figure legends and P values of ≤0.05 were considered significant. All experiments were performed at least twice.

## 3 Results

### 3.1 CYLD expression in eczematous skin lesions of AD patients

The expression of CYLD in different organs and cells under homeostatic conditions has been summarized in “The Human Protein Atlas” (https://www.proteinatlas.org/ENSG00000083799-CYLD). These data show that CYLD is expressed in all organs including the skin with strong protein and mRNA expression in keratinocytes, Langerhans cells, immune cells including T cells, macrophages and B cells but not in melanocytes. (https://doi.org/10.1038/nbt1210-1248). In blood, CYLD is expressed in all leukocyte populations including granulocytes and monocytes (https://www.proteinatlas.org/ENSG00000083799-CYLD/immune+cell). Transcriptome analysis of skin biopsies from 10 AD patients identified CYLD mRNA in T cells, natural killer cells (Fig 1A), and different macrophage subtypes including M2, DCs (Fig 1B). mRNA expression in keratinocytes, fibroblasts, vascular endothelial cells and pericytes appeared weaker but was present as well, while CYLD expression was also detectable in PBMC samples of the same donors including different myeloid cells (Fig 1C, D). Our histological analysis of CYLD expression in AD patients revealed the expression of CYLD in CD3^+^ T cells as well as non-T cells (Fig. 1E). CYLD expression appeared particularly strong in the lesional skin of a patient with clinical signs of superinfected eczema (Fig. 1E, patient AD3). Our further histological analysis confirmed the widespread CYLD expression in the cellular infiltrate and keratinocytes of inflamed AD skin lesions as compared to healthy skin (Suppl. Fig. 1). Thus, CYLD is expressed both in the skin of healthy persons and AD patients with expression in identical cell types including skin macrophages and monocytes in blood.

**Figure 1.**
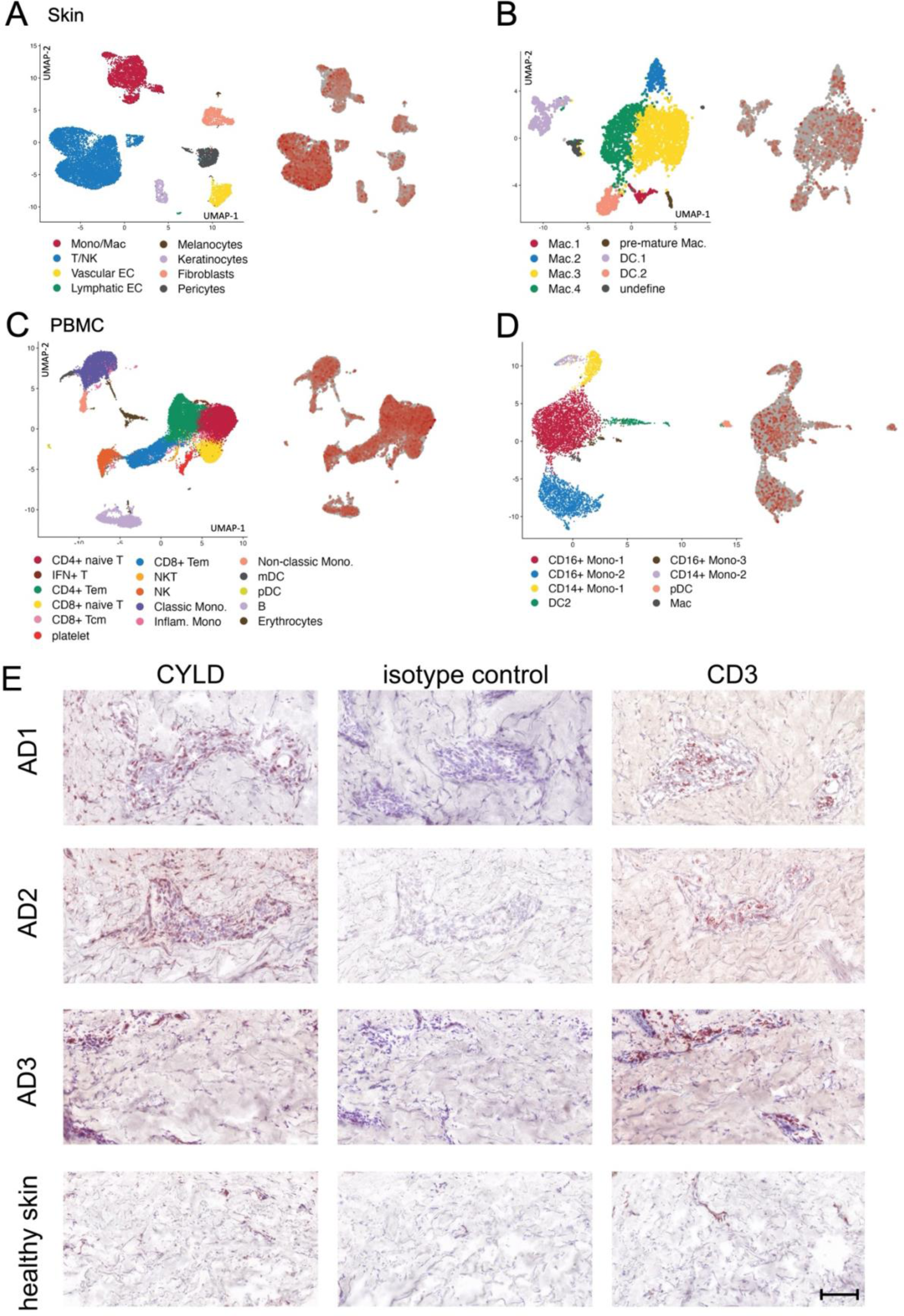
CYLD mRNA and protein expression in AD. (A, B) CYLD mRNA expression in leukocytes (A) and myeloid cell populations (B) isolated from the skin of 10 patients with AD. (C, D) CYLD mRNA expression in PBMC (C) and myeloid cells (D) of the same donors as in (A, B). (E) Exemplary pictures of CYLD protein expression (brown, AEC) in the skin of three patients with AD and healthy control skin. The eczematous lesion of the patient AD3 showed signs of superinfected eczema. Sections were immunostained with a polyclonal rabbit anti-CYLD antibody or a respective isotype control and counterstained with hemalum. Mouse anti-CD3 was applied as a reference. Scale bar = 100µm.

### 3.2 Increased CYLD expression and impaired control of *S. aureus* in monocyte-derived macrophages (MDM) of AD patients

Since AD is characterized by a type 2 inflammation including M2 macrophage polarization (Beck et al., 2022) and activation of the IFN-γ/STAT1 and TLR2/4 pathways synergize in the control of *S. aureus*, we first determined whether IFN-γ/LPS-stimulation improves the control of *S. aureus*-infected macrophages of AD patients to the same extent as in healthy controls. In these experiments, we isolated CD14^+^ monocytes from the blood of AD patients and healthy controls and differentiated these cells into macrophages by M-CSF treatment followed by IFN-γ/LPS treatment. Control of *S. aureus* was significantly impaired in MDM from all investigated AD patients (Fig. 2A). Since CYLD can impair the cell-intrinsic control of intracellular bacteria (Nishanth et al., 2013, Wex et al., 2016), we determined the CYLD expression in the infected MDM. WB analysis showed that the impaired capacity of IFN-γ /LPS-stimulated MDM of AD correlated with an increased CYLD protein expression in these infected cells (Fig. 2B).

**Figure 2.**
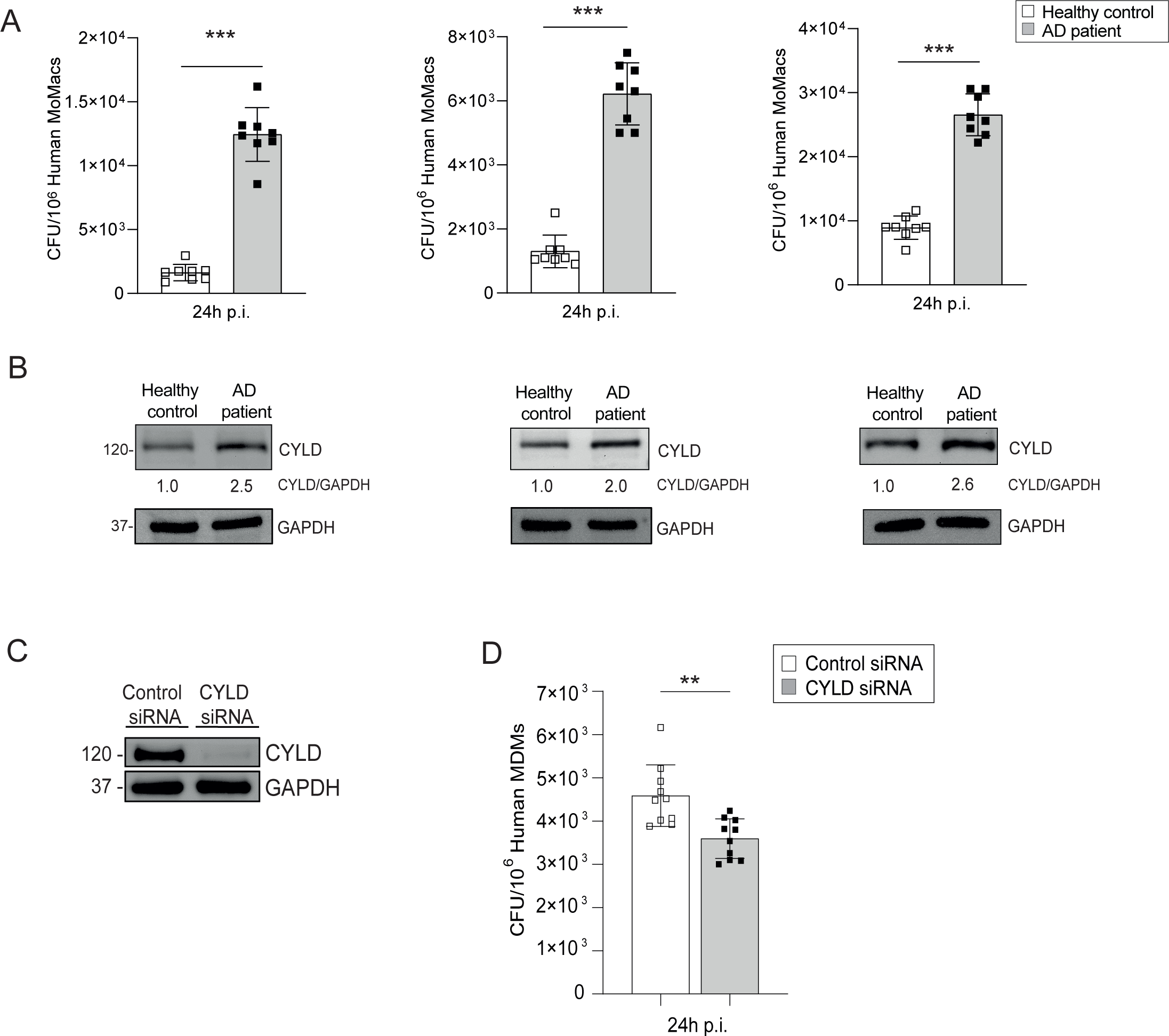
Elevated CYLD expression and impaired control of *S. aureus* by monocyte-derived macrophages from atopic dermatitis patients. CD14^+^ monocytes were isolated from the blood of atopic dermatitis patients and healthy controls using density gradient centrifugation followed by MACS for CD14^+^ cells, thereafter, the monocytes were differentiated into macrophages through M-CSF (50 ng/ml) treatment. (A) The macrophages from healthy control and AD patients were polarized into M1 phenotype by stimulation with IFN-γ (20 ng/ml) and LPS (10 pg/ml) for 24 hours. Cells were harvested and protein lysates were analyzed for CYLD expression by Western blot. GAPDH was used as a loading control. (B) M1 polarized macrophages from atopic dermatitis patients and healthy controls were infected with Staphylococcus aureus (MOI 1) for 24 hours. Thereafter cells were lysed, serial dilutions of the cell lysates were plated on agar plates and the bacterial colonies were enumerated after 24 hours (n=8 per group). (C,D) CD14^+^ monocytes were isolated from buffy coats of healthy controls by density gradient centrifugation and MACS and differentiated into macrophages by M-CSF (50 ng/ml) treatment. Thereafter, cells were transfected with CYLD siRNA or control siRNA, respectively. Twenty-four hours after transfection, monocyte-derived macrophages (MDM) were polarized into M1 by stimulation with IFN-𝛾 (20 ng/ml) and LPS (10 pg/ml) for 24 h. (C) The efficiency of the CYLD knockdown was analyzed 72 h after transfection by WB. (D) After stimulation for 24 h, MDM were infected with *S. aureus* (strain MW2; MOI 1:1). The intracellular bacterial load was determined 24 h p.i. (n=10 per group). Bars represent mean values ±SD (Student’s unpaired t-test, **p<0.01). Data from one of two independent experiments with similar results are shown.

Thus, AD patients are characterized by an overexpression of CYLD in macrophages and impaired control of *S. aureus*. To corroborate the inhibitory function of CYLD on the control of *S. aureus* in human M1-polarized macrophages, we deleted CYLD by siRNA in IFN-γ/LPS-primed (M1) of healthy blood donors (Fig. 2C). Upon infection of CYLD deleted human MDM, the control of *S. aureus* was significantly improved as compared to control siRNA-treated macrophages (Fig. 2D). Thus, CYLD is upregulated in MDM of AD patients and impairs the control of *S. aureus* in M1-polarized MDM demonstrating that CYLD is a macrophage intrinsic inhibitor of the control of *S. aureus*.

### 3.3 CYLD inhibits M1 macrophage polarization, production of anti-bacterial reactive oxygen-species and control of *S. aureus*

To determine the mechanisms how CYLD impairs intrinsic immunity to *S. aureus*, we established *CYLD*-deficient THP1 monocytes by CRISPR/Cas9 (Fig. 3A). Upon M-CSF-mediated differentiation into macrophages and subsequent IFN-γ/LPS-stimulation, *CYLD*-deficient THP1 cells expressed higher levels of CD80, a marker for M1 macrophages, TNF and IL-6 mRNA, two prototypic M1 macrophage cytokines important for the control of *S. aureus* (Fig. 3B). In IFN-γ/LPS-stimulated THP1 macrophages, *CYLD*-deletion significantly improved control of methicillin-sensitive and methicillin-resistant *S. aureus* whereas no differences between the two genotypes were observed for unstimulated macrophages (Fig. 3C, D). A kinetic analysis of the production of anti-bacterial nitric oxide and reactive oxygen species revealed that ROS were significantly increased in infected *CYLD*-deficient THP1 24h p.i. whereas NO levels did not differ (Fig. 3E-G). These data illustrate that CYLD impaired M1 macrophage differentiation, control of *S. aureus* and ROS production.

**Figure 3.**
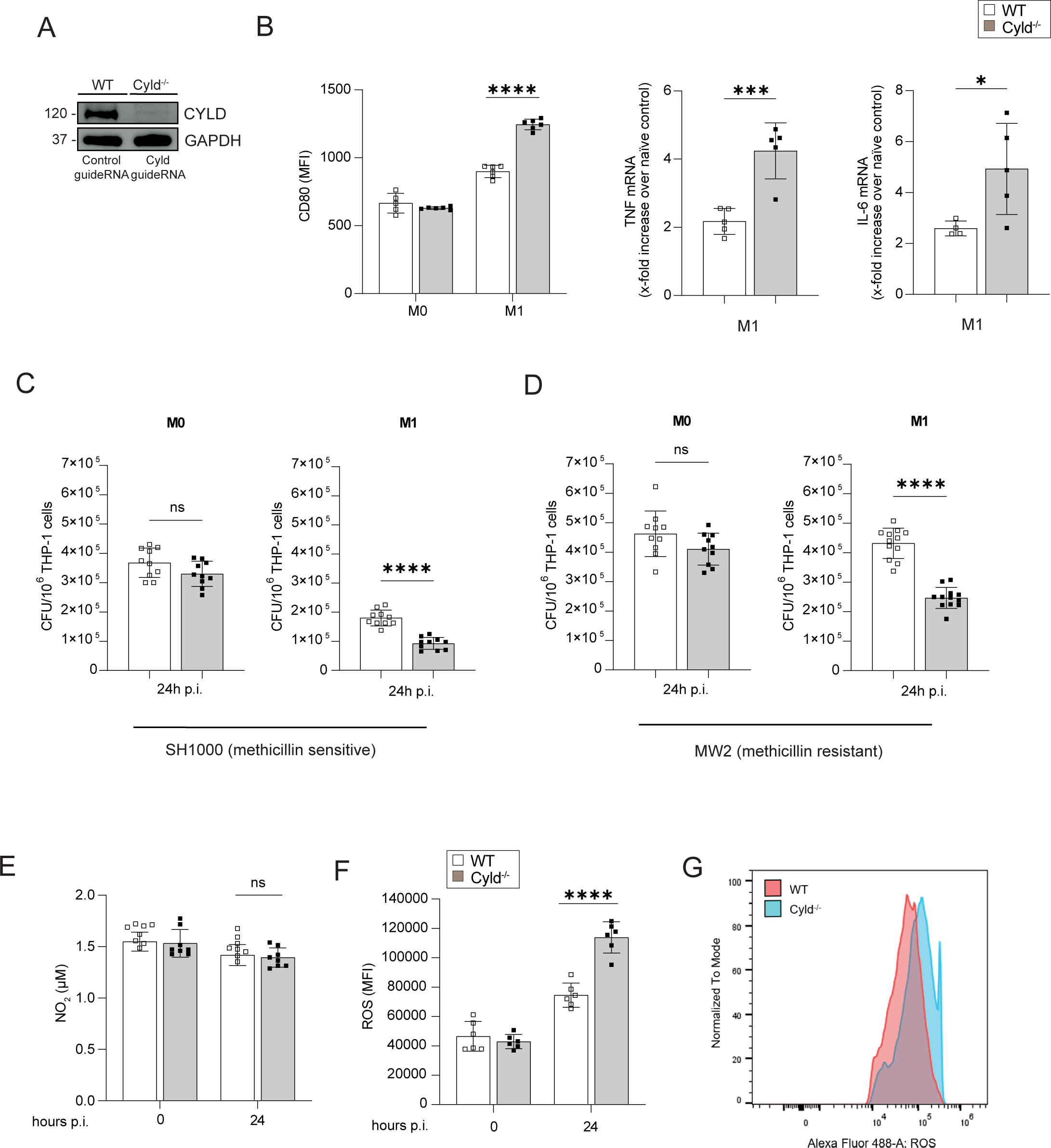
CYLD impairs M1 polarization of macrophages, fosters intracellular control of *S. aureus* in THP-1-derived M1 macrophages and impairs TNF and ROS production in M1 macrophages upon *S. aureus* infection. (A) *CYLD* was deleted in THP1 cells by CRISPR/Cas9 and subsequently differentiated into macrophages through PMA (50 ng/ml) treatment. The efficacy of the CYLD knockout was evaluated via Western blot (WB) analysis. GAPDH was utilized as a loading control. WT and *CYLD*-deficient (*CYLD*^-/-^) THP-1-derived macrophages were polarized into M1 phenotype by stimulation with IFN-𝛾 (20 ng/ml) and LPS (10 pg/ml) for 24 h or left unpolarized (M0). (B) Twenty-four hours after stimulation macrophages were harvested and stained for the M1 marker CD80. Cells were fixed with 4% PFA and the mean fluorescence intensity (MFI) was measured by flow cytometry (n=5 per group). A quantitative RT-PCR (qRT-PCR) analysis of signature cytokines of M1 macrophages TNF and IL-6 mRNA expression was performed 24 h after stimulation (n=6 per group). Changes in gene expression were normalized to HPRT. In (A, B) bars represent mean values ±SD (Student’s unpaired t-test, *p<0.05, ***p<0.001). (C-D) Wild-type (WT) and *CYLD*-deficient (*CYLD*^-/-^) THP-1-derived macrophages were polarized into the M1 phenotype by stimulation with IFN-γ (20 ng/ml) and LPS (10 pg/ml) for 24 hours, followed by infection with Staphylococcus aureus (MOI 1). The intracellular bacterial load was determined in unpolarized M0 and polarized M1 wild-type (WT) and Cyld^-/-^ macrophages 24 hours post-infection (p.i.) with (C) methicillin-sensitive S. aureus (strain SH1000) and (D) methicillin-resistant S. aureus (strain MW2) (n=10 per group). The cells were lysed, serial dilutions of the cell lysates were plated on agar plates and the bacterial colonies were enumerated after 24 hours. In (A and B), the bars represent the mean values ± SD (Student’s unpaired t-test, ***p < 0.001, ****p < 0.0001). The data presented are from one of two independent experiments with similar results. (E-G) Wild-type (WT) and *CYLD*-deficient (*CYLD*^-/-^) THP-1-derived macrophages were polarized into the M1 phenotype by stimulation with IFN-γ (20 ng/ml) and LPS (10 pg/ml) for 24 hours. Thereafter, cells were infected with *S. aureus* (strain MW2; MOI 1:1). (E) The NO_2_ concentration in the supernatant of uninfected (0 h) and infected (24 h p.i.) WT and Cyld^-/-^ macrophages was measured photometrically by the Griess assay (n=8 per group). (F, G) The level of intracellular ROS was determined at (0 h) and 24 h p.i. by flow cytometry using a ROS detection kit (n=6 per group). In (E-F), bars represent mean values ±SD (Student’s unpaired t-test, *p<0.05, ****p<0.0001). Data from one of two independent experiments with similar results are shown.

### 3.4 CYLD inhibits STAT1, Myd88 and NOD2 signaling in *S. aureus*-infected M1 polarized macrophages

Since stimulation with IFN-γ and LPS improved control of *S. aureus* and anti-bacterial activity in *CYLD*-deficient macrophages, we determined the impact of CYLD on (i) IFN-γ-induced STAT1 activation, (ii) the effect of LPS (TLR4) and *S. aureus* (TLR2)-induced MyD88 signaling, and (iii) *S. aureus*-activated NOD2 pathways.

WB analyses showed that CYLD impaired STAT1 phosphorylation at tyrosine 701 and serine 727, which cooperatively lead to the induction of IFN-γ-induced genes (Sadzak et al., 2008; Varinou et al., 2003) (Fig. 4A). Co-immunoprecipitation experiments newly identified that CYLD directly interacted with STAT1 (Fig. 4B, C). STAT1 immunoprecipitation and subsequent WB analysis of K63 ubiquitin revealed that *S. aureus* infection induced increased K63 ubiquitination of STAT1 in *CYLD*-deficient macrophages as compared to WT macrophages at 2h p.i. (Fig. 4D). In agreement with published data in other models (Brummelkamp et al., 2003; Kovalenko et al., 2003; Trompouki et al., 2003; Wex et al, 2016), CYLD also interacted with TRAF6, IKKγ/NEMO and RIPK2 in IFN-γ/LPS-stimulated *S. aureus*-infected THP1 macrophages (Fig. 4E). This was associated with impaired MyD88 and NOD2/RIPK2 signaling leading to reduced downstream NF-κB activation shown by impaired phosphorylation of p65 in *S. aureus*-infected CYLD-competent macrophages (Fig. 4F). Thus, CYLD inhibits simultaneously key pathways leading to reduced activation of the transcription factors NF-κB and STAT1.

**Figure 4.**
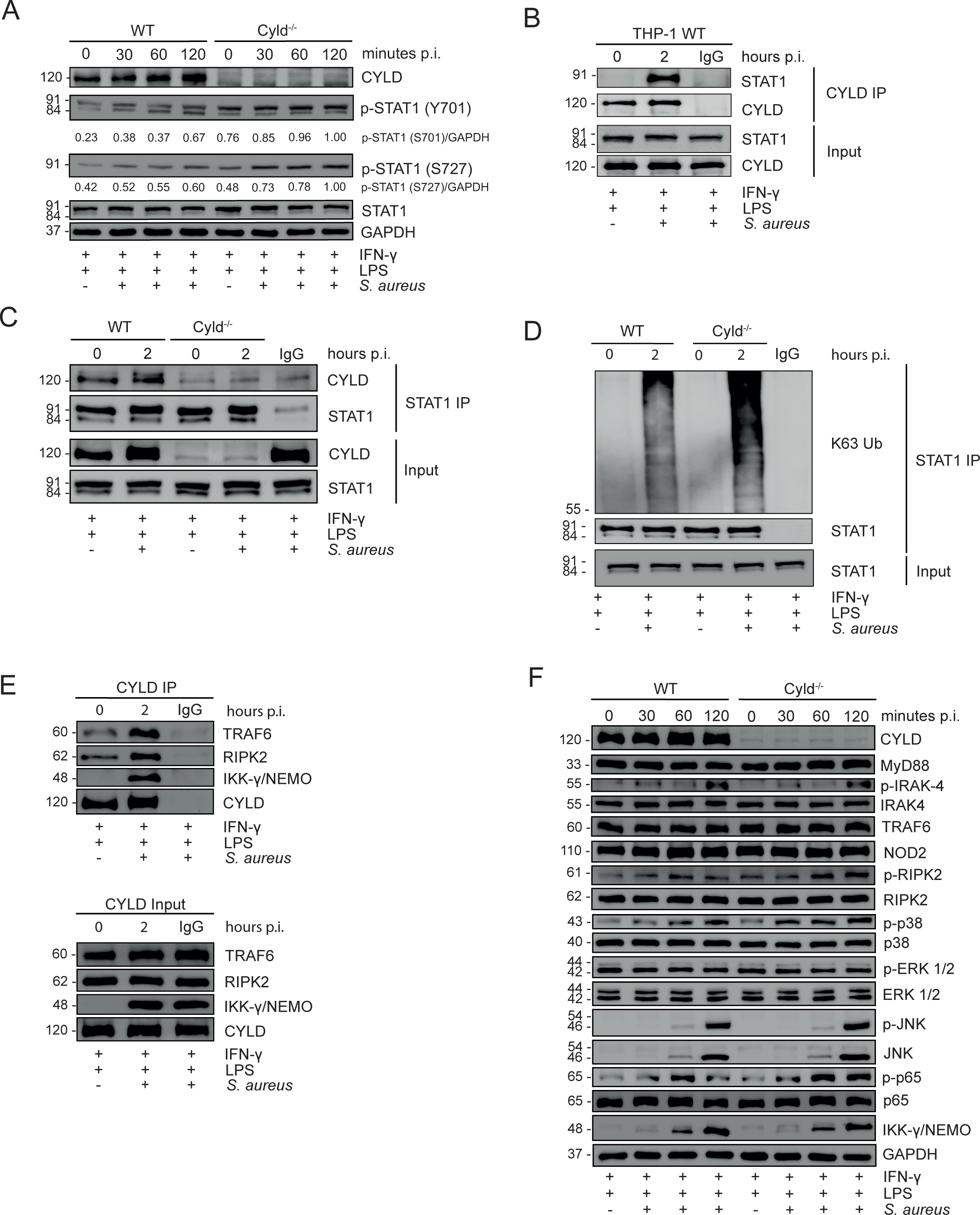
CYLD impairs STAT1, NF-𝜿B and pathways upon *S. aureus* infection. (A-C) WT and *CYLD*^-/-^ THP-1-derived macrophages were polarized into M1 phenotype by stimulation with IFN-𝛾 (20 ng/ml) and LPS (10 pg/ml) for 24 h, Thereafter, cells were infected with *S. aureus* (strain MW2; MOI 5:1) and harvested at 0 (uninfected), 30, 60 and 120 minutes p.i. Proteins were isolated and a western blot analysis of the indicated proteins was performed. GAPDH was used as a loading control. Representative western blots from one of two independent experiments are shown. (A) WT and *CYLD*^-/-^ THP-1-derived macrophages were polarized into M1 by stimulation with IFN-𝛾 (20 ng/ml) and LPS (10 pg/ml) for 24 h. Thereafter, cells were infected with *S. aureus* (strain MW2; MOI 5:1) and harvested after at 0 (uninfected), 30, 60 and 120 minutes p.i. Proteins were isolated and a western blot analysis of the indicated proteins was performed. GAPDH was used as a loading control. Representative western blots from one of two independent experiments are shown. (B) THP1-derived macrophages were polarized into M1 by stimulation with IFN-𝛾 (20 ng/ml) and LPS (10 pg/ml) for 24 h, followed by infection with *S. aureus* (strain MW2; MOI 5:1). Two hours after infection, cells were harvested and proteins were isolated. A co-immunoprecipitation with CYLD antibody was performed and immunoprecipitates were stained for CYLD and STAT1. (C, D) WT and Cyld^-/-^ THP-1-derived macrophages were polarized into M1 by stimulation with IFN-𝛾 (20 ng/ml) and LPS (10 pg/ml) for 24 h followed by infection with *S. aureus* (strain MW2; MOI 5:1). Two hours after infection, cells were harvested and protein lysates were co-immunoprecipitated with STAT1 antibody. Western blot analysis for CYLD (C), STAT1 (C, D) and K63-linked polyubiquitin (D) was performed. In (B-D), representative western blots from one of three independent experiments are shown. (E) WT and *CYLD*^-/-^ THP-1-derived macrophages were polarized into M1 by stimulation with IFN-𝛾 (20 ng/ml) and LPS (10 pg/ml) for 24 h followed by infection with *S. aureus* (strain MW2; MOI 5:1). Two hours after infection, cells were harvested and protein lysates were co-immunoprecipitated with CYLD antibody. Western blot analysis for CYLD, RIPK2 and IKKγ/NEMO was performed. (F)WT and *CYLD*^-/-^ THP-1-derived macrophages were polarized into M1 phenotype by stimulation with IFN-𝛾 (20 ng/ml) and LPS (10 pg/ml) for 24 h, Thereafter, cells were infected with *S. aureus* (strain MW2; MOI 5:1) and harvested at 0 (uninfected), 30, 60 and 120 minutes p.i. Proteins were isolated and a western blot analysis of the indicated proteins was performed. GAPDH was used as a loading control.

### 3.5 The enhanced control of intracellular *S. aureus* in *CYLD*-deficient THP-1-derived M1 macrophages is NF-**𝜿**B- and STAT1-dependent

To determine the functional importance of CYLD regulated NF-κB and STAT1 activation for the control of *S. aureus*, we stimulated THP1 macrophages with IFN-γ/LPS and treated the cells prior to infection with the IKK VII inhibitor to inhibit activation of NF-κB and with STAT1 siRNA to inhibit STAT1, respectively. Inhibition of NF-κB activation and STAT1, respectively, resulted in a strong increase of CFU of *S. aureus* in both WT and *CYLD*-deficient THP1 macrophages and both treatments abolished the differences between the two genotypes (Fig. 5A, B). Thus, the inhibition of both STAT1 and NF-κB by CYLD is both critical for the control of *S. aureus* in macrophages.

**Figure 5.**
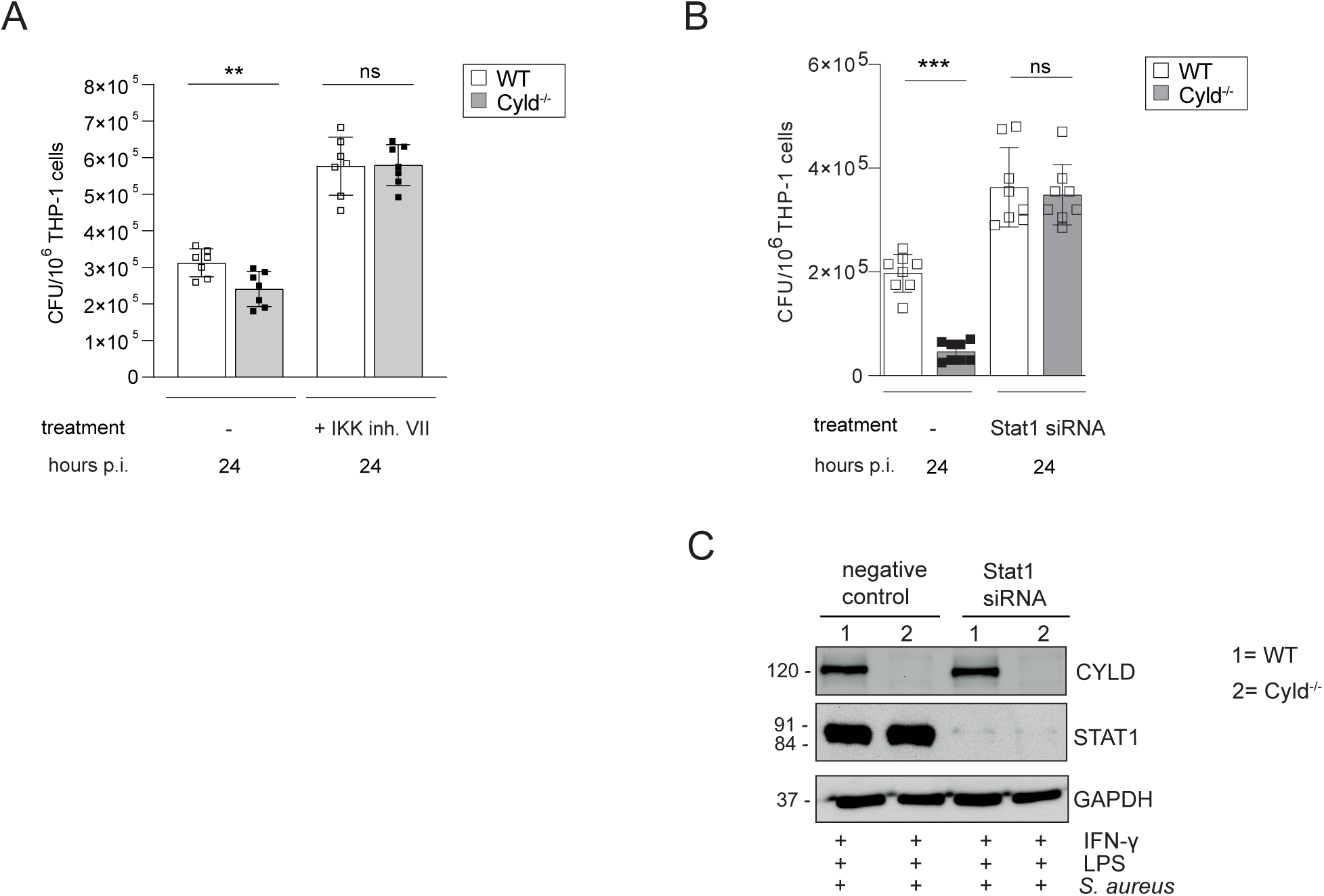
The improved control of intracellular *S. aureus* in Cyld-deficient THP-1-derived M1 macrophages is NF-𝜿B- and STAT1-dependent. WT and Cyld^-/-^ THP-1-derived macrophages were polarized into M1 phenotype by stimulation with IFN-𝛾 (20 ng/ml) and LPS (10 pg/ml) for 24 h. In parallel, cells were either left untreated or treated with IKK inhibitor VII (1 𝜇M), thereafter, the cells were infected with *S. aureus* (strain MW2, MOI 1:1). The inhibitor treatment was continued during and after infection. Twenty-four hours p.i., the intracellular bacterial load of the different groups of MRSA-infected WT and Cyld^-/-^ macrophages was determined (n=6 per group). (B) For STAT1 knockdown, STAT1 siRNA was added to WT and *CYLD*^-/-^ macrophages 24 h prior to addition of IFN-𝛾 (20 ng/ml) and LPS (10 pg/ml) followed by infection with *S. aureus* strain MW2 (MOI 1:1). The intracellular bacterial load was determined 24h p.i. (C) Efficiency of STAT1 knockdown was validated using western blotting. Bars represent mean values ±SD (Student’s unpaired t-test, **p<0.01). Data from one of two independent experiments with similar results are shown.

3.6 *Cyld*-deficient mice are protected from *S. aureus* infection

The data presented identify that CYLD is increasingly expressed in macrophages of AD patients and inhibits the control of *S. aureus* in human macrophages. To further validate an inhibitory function of CYLD in *S. aureus* infection and to evaluate whether systemic CYLD inhibition might by a therapeutic option to ameliorate *S. aureus* infection, we infected *Cyld*-deficient and WT mice with *S. aureus*. *Cyld*-deficient mice had a significantly reduced weight loss (Fig. 6A) and an improved pathogen control in liver and kidney at the acute and chronic stage of infection (Fig. 6B, C). This, qualifies CYLD as relevant therapeutic target to ameliorate *S. aureus* infections.

**FIGURE 6.**
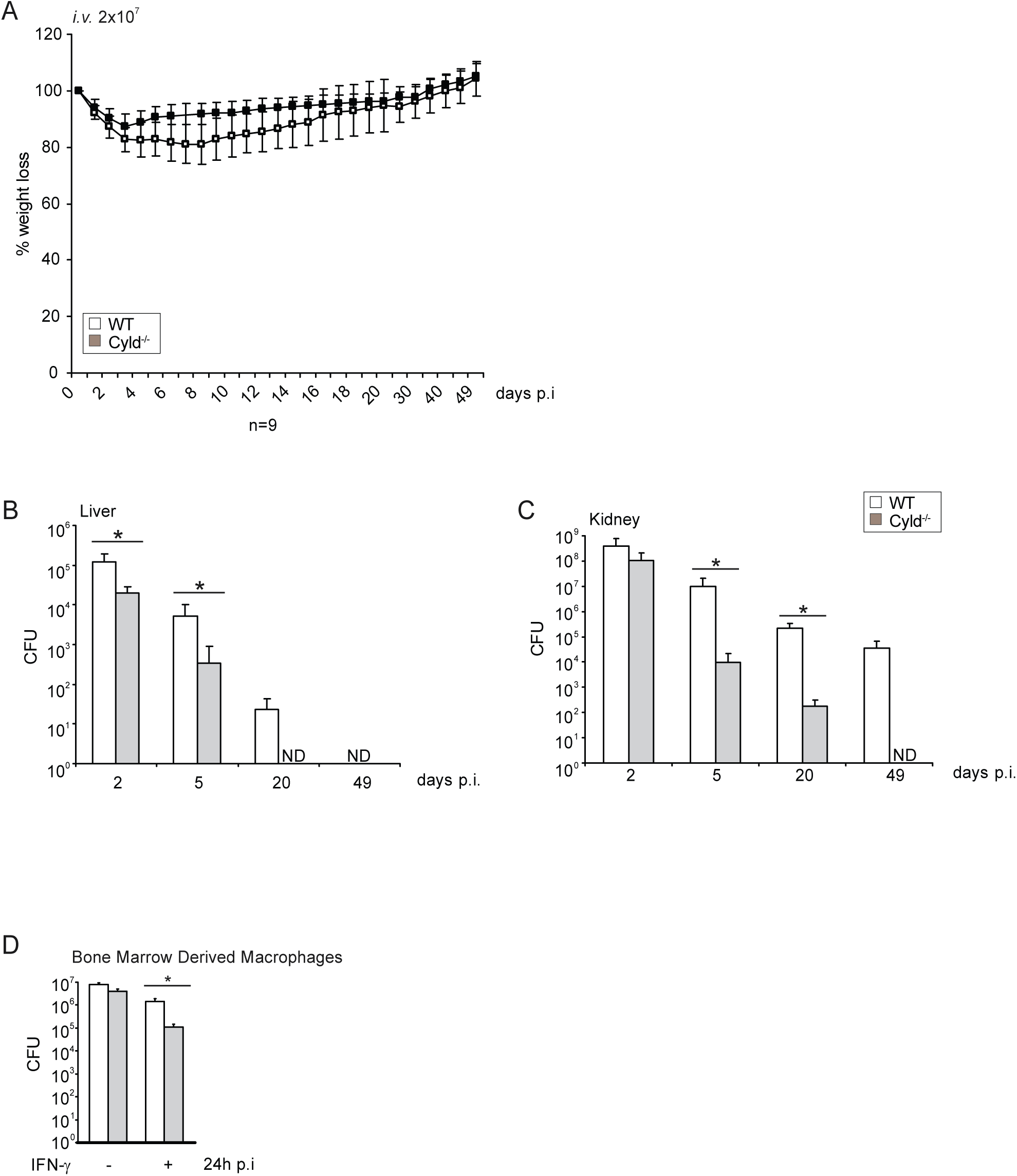
CYLD-deficient mice are protected from *S. aureus* infection. (A-C) C57BL WT and Cyld^-/-^ mice (8-10 weeks old) were intravenously infected with 2x10^7^ *S. aureus* (strain SH1000). (A) Weight of mice was monitored daily up to day 50 p.i. The weight on the day of infection was defined as 100% and the percentage of differences in weight compared to the 100% are shown (n=6-8 mice per group). (B, C) Mice were sacrificed at the indicated timepoints p.i. and 10-fold serial dilutions of organ homogenates were plated on agar plates. After 24 h, colonies were counted microscopically and the colony forming units (CFU) were calculated for liver (B) and kidney (C) (n=6 per timepoint and group). (D) Triplicates of bone-marrow-derived macrophages (BMDM) were treated with IFN-𝛾 (100 U/ml) for 6 h before infection with *S. aureus* in a gentamicin protection assay (MOI 5:1) for 24 h. Thereafter, cells were lysed and CFU were determined. In (A-E) the mean values ±SD are shown (Student’s unpaired t-test, *p<0.05). Data from one of two independent experiments with similar results are shown.

## 4 Discussion

Macrophages play a fundamental role in the control of local and systemic infections with *S. aureus* (Pidwill et al., 2021; Surewaard et al., 2016; Vajjala et al., 2016). In murine *S. aureus* skin infection M1-polarized macrophages contribute to the control of the infection, whereas M2 polarized macrophages are associated with impaired killing of *S. aureus* and spread of the pathogen (Asai et al., 2010). In AD, the cutaneous inflammatory lesions are characterized by a type 2 inflammatory milieu including M2 polarized macrophages infiltrating the diseased skin (Hashimoto et al., 2023; Kasrai and Werfel, 2013; Zhang et al., 2023). A shift to M1 macrophages would reduce *S. aureus* colonization and infections in AD patients. However, this study illustrates that IFN-γ/LPS-stimulated MDM of AD patients express increased amount of CYLD, and have an impaired capacity to kill *S. aureus*. The functional importance of CYLD is shown in THP1 macrophages which only incompletely shift to an M1 phenotype as illustrated by reduced CD80 expression and TNF production and by the improved control of *S. aureus* in *CYLD*-deficient IFN-γ/LPS activated MDM of healthy blood donors and THP1 cells.

The present study newly identifies that CYLD binds to STAT1 upon *S. aureus* infection of IFNγ/LPS-primed MDM. This interaction resulted in a CYLD-dependent reduction of K63 polyubiquitination of STAT1 and an impaired STAT1 phosphorylation. CYLD inhibited both the phosphorylation at STATY701 which is important for nuclear accumulation of the STAT1 transcription factor, and the subsequent nuclear phosphorylation at STAT1S727 which is required for gene transcription (Sadzak et al., 2008; Varinou et al., 2003). The functional importance of STAT1 for the control of *S. aureus* was illustrated by the impaired control of *S. aureus* in IFN-γ/LPS-stimulated MDM upon siRNA-mediated inhibition of STAT1.

The induction of an anti-staphylococcal function of macrophages also requires the engagement of pathogen pattern recognition receptors leading to the activation of NF-κB (Pidwill et al., 2021). In the present study, the pre-stimulation with the TLR4 agonist LPS and the subsequent *S. aureus* infection activating TLR2 and NOD2 by lipoteichoic acid and muramyl dipeptide, respectively, led to the activation of NF-κB and NF-κB-dependent production of anti-bacterial ROS. Production of ROS is important and has a superior role in comparison to NO to control of *S. aureus* infections (Hashimoto et al., 2006; Pizzolla et al., 2012; Schäffler et al., 2014; Surewaard et al., 2016; Takeuchi et al, 1999; Takeuchi et al., 2000). In agreement with previous studies, we detected an interaction of CYLD with RIPK2, TRAF6 and IKKγ/NEMO, which are critical signaling molecules in the NOD2 and TLR2/4 pathways leading to NF-κB activation (Brummelkamp et al., 2003; Kovalenko et al., 2003; Trompouki et al., 2003; Wex et al, 2016). CYLD inhibits the activation of these signaling molecules by its K63 deubiquitinating activity. The key role of CYLD-mediated NF-κB inhibition as a critical factor impairing control of *S. aureus* in macrophages is proven by the strong increase and abolishment of differences in CFU between CYLD-competent and -deficient macrophages. Since numbers of S. aureus were identical in both NF-κB inhibited and also in STAT1 siRNA treated WT and *CYLD*^-/-^ macrophages, the two pathways determine independently from each other the control of *S. aureus* and cannot compensate for each other.

Of note, infection with *S. aureus* led to a CYLD-independent and equal activation of MAP kinases including c-Jun-N-terminal kinase (JNK), p38 and ERK in CYLD-competent and -deficient MDM. Previously we have shown that the inhibition of RIPK2 by CYLD impairs the activation of MAP kinases and ERK1/2-induced autophagy in *Listeria monocytogenes* infected macrophages (Wex et al., 2016). In the present study, MAP kinases, in particular JNK, were equally activated in CYLD-deficient and -competent MDM. This difference may be explained by the manipulation of MAP kinases by *S. aureus*, which may undermine JNK function by a yet unresolved mechanism to induce its persistence in macrophages (Watanabe et al., 2007).

Our data show that combined PRR-mediated activation of NF-κB and IFN-γ-driven STAT1 activation augments the anti-staphylococcal activity of macrophages. Unleashing the CYLD break on NF-κB and STAT1 activation further enhances the control of *S. aureus* of M1 polarized macrophages. The in vivo data illustrating that *Cyld*^-/-^ mice have an improved course and control of *S. aureus* in acute and chronic systemic infection further corroborate that CYLD is a potential therapeutic target to reduce *S. aureus* colonization and infection.

## 6 Author Contributions

CS and ST performed experiments and analyzed data. KH performed experiments, analyzed data and wrote the manuscript. GN and LN planed and performed experiments, analyzed data and wrote the manuscript. TW and DS planed experiments, analyzed data and wrote the manuscript.

## 8 Acknowledgments

The authors thank Birgit Brenneke, Kerstin Ellrott, and Gabriele Begemann for expert technical assistance. Further we thank all volunteers for their contribution to this project.

## 10 Data Availability Statement

Publicly available datasets were analyzed in this study. This data can be found here: https://ega-archive.org/EGAD00001010106.

## 12 Supplementary Material

**Supplementary Figure 1:**
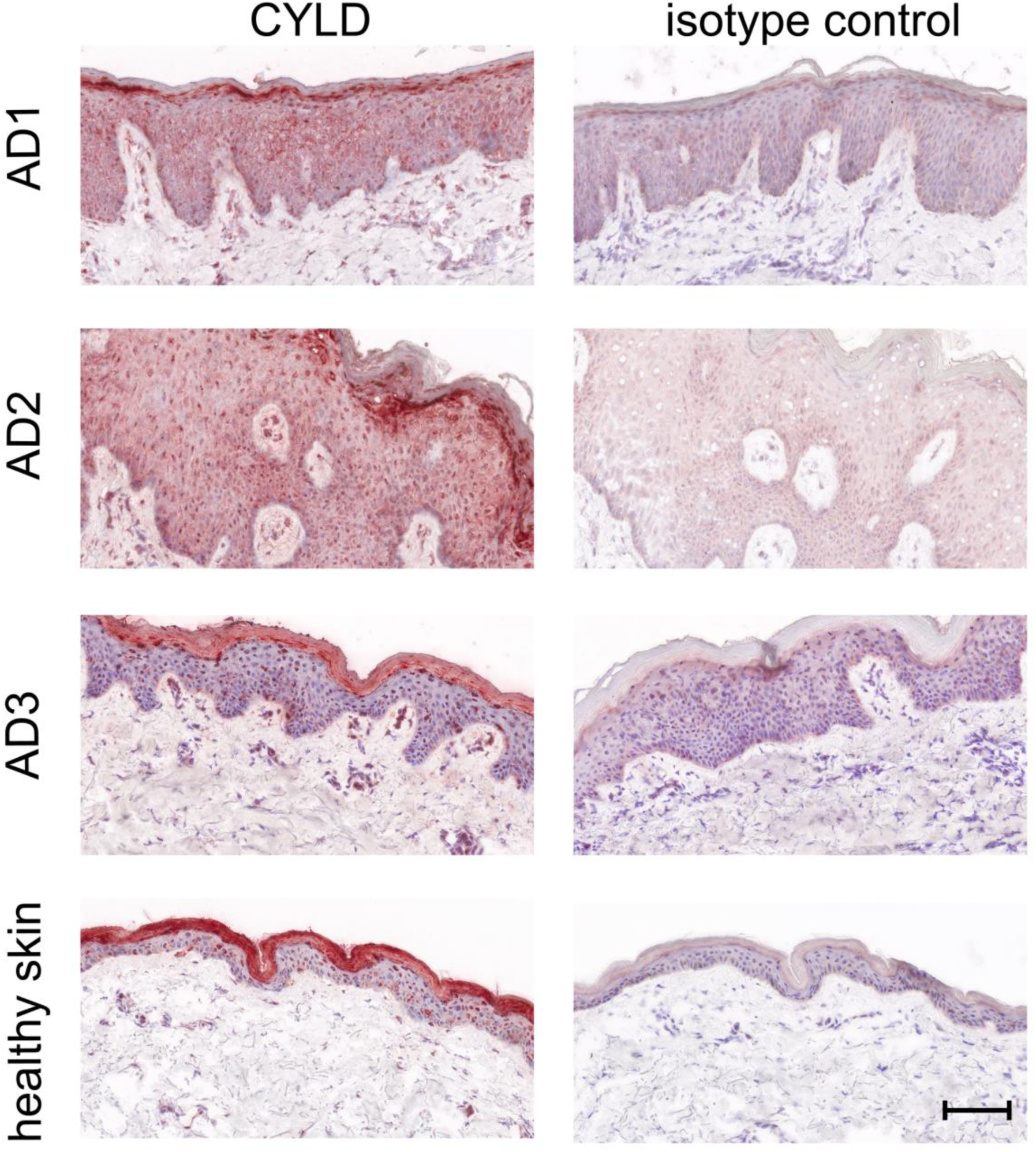
Increased CYLD protein expression in the epidermal skin of atopic dermatitis patients. Exemplary pictures of CYLD protein expression (brown, AEC) in the skin of three patients with AD and one healthy control. Sections were immunostained with a polyclonal rabbit anti-CYLD antibody or a respective isotype control and counterstained with hemalum. Scale bar = 100µm.

